# *Tetrahymena* Poc5 is a transient basal body component that is important for basal body maturation

**DOI:** 10.1101/812503

**Authors:** Westley Heydeck, Brian A. Bayless, Alexander J. Stemm-Wolf, Eileen T. O’Toole, Courtney Ozzello, Marina Nguyen, Mark Winey

**Author notes:** CORRESPONDING AUTHOR: Mark Winey, Department of Molecular and Cellular Biology, (530)752-6778.

## Abstract

Basal bodies (BBs) are microtubule-based organelles that template and stabilize cilia at the cell surface. Centrins ubiquitously associate with BBs and function in BB assembly, maturation, and stability. Human POC5 (hPOC5) is a highly conserved centrin-binding protein that binds centrins through Sfi1p-like repeats and is required for building full-length, mature centrioles. Here, we use the BB-rich cytoskeleton of *Tetrahymena thermophila* to characterize Poc5 BB functions. *Tetrahymena* Poc5 (TtPoc5) uniquely incorporates into assembling BBs and is then removed from mature BBs prior to ciliogenesis. Complete genomic knockout of *TtPOC5* leads to a significantly increased production of BBs yet a markedly reduced ciliary density, both of which are rescued by reintroduction of TtPoc5. A second *Tetrahymena POC5*-like gene, *SFR1*, is similarly implicated in modulating BB production. When *TtPOC5* and *SFR1* are co-deleted, cell viability is compromised, and levels of BB overproduction are exacerbated. Overproduced BBs display defective transition zone formation and a diminished capacity for ciliogenesis. This study uncovers a requirement for Poc5 in building mature BBs, providing a possible functional link between *hPOC5* mutations and impaired cilia.

**SUMMARY STATEMENT:** Loss of *Tetrahymena thermophila* Poc5 reveals an important role for this centrin-binding protein in basal body maturation, which also impacts basal body production and ciliogenesis.

## INTRODUCTION

Centrioles and basal bodies (BBs) are evolutionarily ancient, microtubule-based organelles that nucleate and organize microtubules into specific arrangements for fundamental cellular processes (Carvalho-Santos et al., 2011; Hodges et al., 2010; Kobayashi and Dynlacht, 2011). During cell division, microtubule arrays emanating from centriole-comprised centrosomes function to assemble and organize the mitotic spindle for cellular control of chromosome segregation. BBs are structurally similar to centrioles, and in some cell types centrioles and BBs are interconverted as a function of the cell cycle; however, BBs are distinctly required for templating, orienting, and anchoring cilia and flagella at the cell surface. Depending on the cell type, cilia and flagella have central roles in diverse mechanical and sensory functions, including generating directional fluid flow for cell motility and normal left-right axis determination, as well as coordinating several important signal transduction pathways (Ishikawa and Marshall, 2011; Nachury and Mick, 2019; Nonaka et al., 1998). In humans, cilia are present on almost all cell types and ciliary dysfunction is associated with complex disorders known as ciliopathies, which display a broad spectrum of clinical features reflective of the ubiquity and functional diversity of cilia (Badano et al., 2006; Waters and Beales, 2011). Further, investigating ciliopathies has solidified the importance of the cilium-associated BB for ciliary function, as many genetic mutations underlying ciliopathies encode defective BB proteins as well as proteins that reside and function in both BBs and centrioles (Reiter and Leroux, 2017).

The characteristic cylindrical structure of centrioles and BBs is primarily composed of a nine-fold, radially symmetric arrangement of specialized triplet microtubules. The definitive nine-fold internal symmetry is imparted during early assembly by the proximal cartwheel structure, which consists of nine spokes radiating from a central hub, with the tip of each spoke typically containing a set of triplet microtubules (Guichard et al., 2018; Hirono, 2014; Kilburn et al., 2007). As centrioles and BBs mature, these structures become polarized with an apparent transition from triplet-to-doublet microtubules occurring at the distal end, as well as the variable acquisition of subdistal and distal appendages (Pearson, 2014; Tanos et al., 2013). Doublet microtubules of the ciliary axoneme originate from the distal end of a mature BB, in a region known as the transition zone (TZ), which is an important subdomain for initiating the compartmentalization of the cilium during early assembly and maintaining a distinct ciliary protein composition (Garcia-Gonzalo and Reiter, 2017; Gonçalves and Pelletier, 2017). Thus, a mature TZ is required for ciliary assembly and maintained ciliary function, and a significant number of ciliopathy-associated proteins localize to and function in the TZ (Reiter and Leroux, 2017).

Centriole and BB duplication is a tightly regulated process generally coupled with the cell cycle, to ensure that constant numbers of these structures are maintained after cell division for specific cellular requirements (Fırat-Karalar and Stearns, 2014; Gönczy, 2012; Pearson and Winey, 2009). To maintain constant centriole and BB numbers, duplication occurs only once per cell cycle and new assembly is typically limited to only one site on a preexisting structure. This assembly process differs in multiciliated cells, where the requisite near-simultaneous assembly of BBs occurs during differentiation and proceeds by multiple BBs forming on a preexisting, template BB as well as by *de novo* assembly around a deuterosome (Brooks and Wallingford, 2014; Dawe et al., 2007; Yan et al., 2016). Notably, numerous studies in diverse model systems have elucidated a strong evolutionary conservation of core molecules that regulate duplication of these structures, including molecular assembly factors with shared functions across assembly pathways (Rodrigues-Martins et al., 2008; Strnad and Gönczy, 2008; Vladar and Stearns, 2007).

Centrins are small calcium-binding proteins that are widely conserved in microtubule-organizing centers including the yeast spindle pole body (yeast centrosome), centrioles, and BBs, where enrichment in centrioles and BBs is found at both the site of new assembly and in the distal portion of these structures (Baum et al., 1986; Kilburn et al., 2007; Laoukili et al., 2000; Paoletti et al., 1996a; Alexander J Stemm-Wolf et al., 2005; Vonderfecht et al., 2012). While the role of centrins in centrosome duplication is less understood, centrins have key functions that contribute to BB duplication and maintenance, including roles in BB assembly, maturation, separation, and stability (Koblenz et al., 2003; Ruiz et al., 2005; Alexander J Stemm-Wolf et al., 2005; Vonderfecht et al., 2011, 2012). Further, defective ciliary assembly and function, as well as an array of ciliopathy-like phenotypes, have been observed in mouse and zebrafish studies of centrin depletion (Delaval et al., 2011; Ying et al., 2019, 2014). Notably, centrins also perform non-centrosomal functions, including roles in DNA damage repair, mRNA export, and fibroblast growth factor-mediated signaling (Dantas et al., 2011; Nishi et al., 2005; Shi et al., 2015).

An interaction between centrin and centrosomal Sfi1p is required for duplication of the yeast spindle pole body and in mammalian cells, the mammalian ortholog of SFI1 promotes centriole duplication (Kilmartin, 2003; Kodani et al., 2019). Structural studies of Sfi1 uncovered centrin-binding repeats (CBRs) that each contain a conserved sequence motif, Ax7LLx3F/Lx2WK/R, that directly binds the C-terminus of centrin (Kilmartin, 2003; Li et al., 2006; Martinez-Sanz et al., 2010, 2006). This conserved motif enabled the identification of centrin-binding proteins across eukaryotes that contain centrioles and/or BBs, including five in humans (Azimzadeh et al., 2009; Eisen et al., 2006; Gogendeau et al., 2007; Heydeck et al., 2016; Kilmartin, 2003; Stemm-Wolf et al., 2013). The centrin-binding protein, Poc5, was identified previously in the *Chlamydomonas reinhardtii* centriole proteome and characterized as an ancestral, core centriolar protein based on broad evolutionary conservation among eukaryotes (Hodges et al., 2010; Keller et al., 2009). Further, human POC5 (hPOC5) was found to be enriched in the distal portion of human centrioles, where it has an essential role in centriole elongation and maturation (Azimzadeh et al., 2009). This role for hPOC5 is notable because molecular mechanisms that contribute to building a full-length centriole/BB remain poorly understood, especially compared with the mechanistic and molecular understanding of early assembly (Chang et al., 2016; Chen et al., 2017; Comartin et al., 2013; Keller et al., 2009; Schmidt et al., 2009). Furthermore, a truncating mutation in *hPOC5* was recently implicated in an inherited form of retinal degeneration, retinitis pigmentosa, characterized by progressive loss of photoreceptors, and Poc5 was found to colocalize with centrin in the connecting cilium of zebrafish photoreceptors, where it is important for normal retinal development and function (Weisz Hubshman et al., 2018; Wheway et al., 2014). Additionally, *hPOC5* mutations associated with adolescent idiopathic scoliosis lead to mislocalization of hPOC5, impaired cell cycle progression, and shorter cilia (Hassan et al., 2019).

The ciliate, *Tetrahymena thermophila*, has a microtubule-based cytoskeleton containing roughly 750 BBs organized along cortical rows and within the feeding structure of the cell, the oral apparatus (Bayless et al., 2015). Previous *Tetrahymena* studies have provided fundamental insight on the multiple BB functions of centrins (Kilburn et al., 2007; Alexander J Stemm-Wolf et al., 2005; Vonderfecht et al., 2011, 2012). Further, members of the large family of 13 centrin-binding proteins in *Tetrahymena* (Sfr1-13) each localize distinctly to subsets of cortical row and/or oral apparatus BBs, as well as to specific regions within BBs overlapping with known patterns of centrin localization (Heydeck et al., 2016; Stemm-Wolf et al., 2013). Also, centrin-binding proteins have diverse roles in BBs, where Sfr1 and Sfr13 function in modulating BB production and in separating/stabilizing BBs, respectively. Together, these findings suggest that centrin-binding proteins may collectively provide the precise spatiotemporal coordination necessary for centrin to perform multiple BB functions. Despite the strong evolutionary conservation of Poc5 in centrioles and BBs, this study provides the first functional characterization of Poc5 in BBs. This investigation reveals a dynamic BB localization pattern for *Tetrahymena* Poc5 (TtPoc5) and an essential role for Poc5 in building a mature BB capable of nucleating a cilium.

## RESULTS

### Identification of TtPoc5 through evolutionarily conserved Poc5 domains

The hallmark feature of centriole and BB-associated centrin-binding proteins is the presence of a variable number of CBRs that each contain a conserved sequence motif, Ax7LLx3F/Lx2WK/R (Kilmartin, 2003; Li et al., 2006). Despite the expanded number of centrin-binding proteins in *Tetrahymena*, mining the *Tetrahymena* genome using solely this conserved sequence motif within CBRs was not a sufficient method for identifying TtPoc5. This was primarily due to limited sequence conservation between centrin-binding proteins, as exemplified by moderate overall sequence homology (mean 16% identity, 29% similarity) even within the Poc5 family (Azimzadeh et al., 2009). Despite limited overall sequence homology, Poc5 orthologs can be differentiated from other centrin-binding proteins because they contain a 21 amino acid (AA) signature sequence motif, known as the Poc5 box, that is both highly conserved (mean 57% identity, 81% similarity) and uniquely found only in Poc5 orthologs (Azimzadeh et al., 2009). Using the full protein sequence of hPOC5 to search the *Tetrahymena* genome, a Poc5 ortholog was identified that shares characteristic Poc5 domain organization and sequence homology with hPOC5 (overall identities, 68/291 [23%]; positives, 135/291 [46%]) (Fig. 1A). Similar to hPOC5, TtPoc5 (TTHERM_00079160) has three CBRs organized as an isolated N-terminal CBR (CBR1) and two CBRs (CBR2,3) in tandem (Fig. 1A,B) (Azimzadeh et al., 2009). Notably, the defining residues of the conserved sequence motif between the three CBRs of hPOC5 and TtPoc5 are primarily either conserved or their hydrophobicity is retained despite AA differences, suggesting that the functional centrin-binding property of TtPoc5 CBRs is likely intact (denoted by asterisks in Fig. 1B, highlighted in Fig. 1C). Additionally, TtPoc5 contains the characteristic Poc5 box, which exhibits a high degree of shared sequence homology with hPOC5 (overall identities, 15/21 [71%]; positives, 17/21 [81%]) (Fig. 1B) and with select Poc5 orthologs (Fig. 1D) (Azimzadeh et al., 2009).

**Fig. 1.**
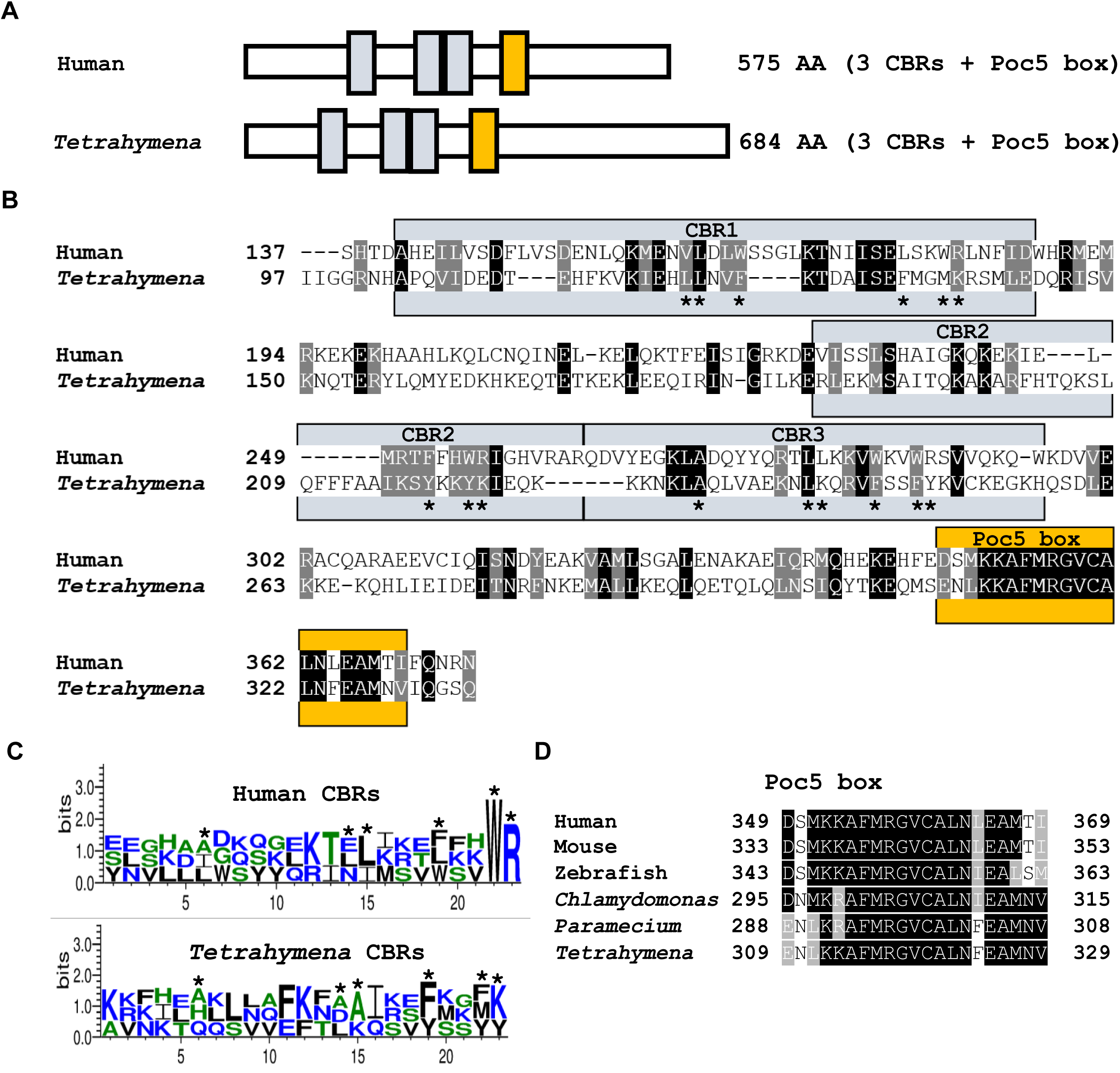
*Tetrahymena* Poc5 (TtPoc5) contains evolutionarily conserved Poc5 domains. **(A)** Schematics showing conserved organization of centrin-binding repeats (CBRs; gray boxes) and the Poc5 box (yellow box) in hPOC5 and TtPoc5. **(B)** Sequence alignment within region containing an isolated CBR (CBR1), tandem CBRs (CBR2,3), and the Poc5 box. Identical residues shaded in black and similar residues shaded in gray. Asterisks denote conserved centrin-binding sequence motif (Ax7LLx3F/Lx2WK/R) residues. **(C)** Sequence logos show AA composition within 18 AA sequence motifs (positions 6-23) of hPOC5 and TtPoc5 CBRs plus five upstream AAs. Black residues indicate high proportion of hydrophobic AAs at conserved positions (marked with asterisks). Hydrophilic residues are indicated in blue and neutral residues are indicated in green. **(D)** Multiple sequence alignment of the highly conserved 21 AA Poc5 box across select Poc5 orthologs.

### TtPoc5 localizes exclusively to assembling BBs

*Tetrahymena* cells have a highly organized cytoskeleton with hundreds of BBs maintained along cortical rows as well in the BB-comprised oral apparatus (Bayless et al., 2015). To determine whether TtPoc5 localizes to BBs, *Tetrahymena* cells containing endogenously tagged Poc5-GFP and the BB-specific marker, Poc1-mCherry, were imaged for colocalization (Heydeck et al., 2016; Pearson et al., 2009; Stemm-Wolf et al., 2013). Endogenous TtPoc5 only colocalized with Poc1 in a subset of cortical row BBs (highlighted with boxes in Fig. 2A) and was notably absent from the oral apparatus (arrowheads in Fig. 2A). Notably, live imaging of *Tetrahymena* cells in the GFP channel captured autofluorescence in large cell vacuoles that was not reflective of Poc5-GFP signal (Fig. 2A).

**Fig. 2.**
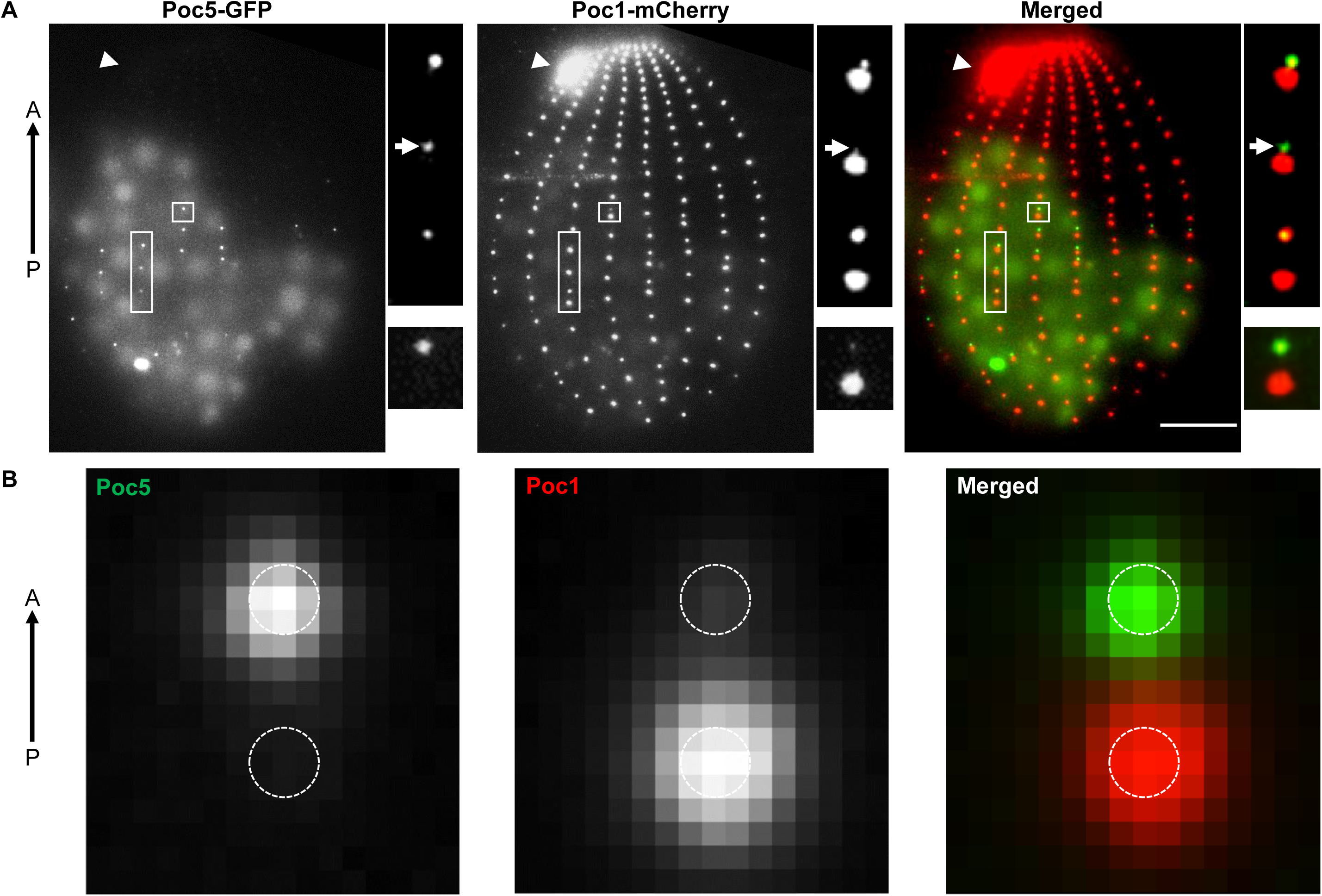
Endogenously tagged Poc5-GFP localizes to assembling BBs. **(A)** Live-cell imaging of C-terminally tagged Poc5-GFP relative to Poc1-mCherry (BB marker) with the anterior(A)-posterior(P) axis indicated with an arrow. Poc5-GFP localizes to a subset of cortical row BBs (highlighted with boxes) but is not detected in the mature oral apparatus (labeled with arrowheads). Scale bar: 10µm. Upper right panels are representative regions of cortical rows containing three BB pairs. Poc5-GFP localizes to the anterior (assembling) BBs and is absent in the posterior (mature) BBs. Poc5 incorporation can precede that of Poc1 (labeled with arrows). Small 1.1 µm x 1.2 µm panels show representative BB pair for image averaging in **(B). (B)** Image averaging of Poc5-GFP and Poc1-mCherry signals across 58 cortical row BB pairs reveals Poc5-GFP localization exclusively at assembling BBs. The BB scaffolds are approximated with 200 nm diameter dashed circles based on the center of the Poc1-mCherry signal, reflecting the average diameter of a *Tetrahymena* BB.

To dissect further the TtPoc5 BB localization pattern, analysis focused on characterizing the Poc5-GFP signal relative to Poc1-mCherry in cortical row BBs. In *Tetrahymena*, new BBs assemble anteriorly from existing, mature BBs in cortical rows and can be identified in closely adjoining BB pairs (Bayless et al., 2015). Endogenously tagged Poc1-mCherry was used for this study because it slowly incorporates into assembling BBs and gradually accumulates during BB maturation, therefore differentiating assembling from mature BBs (Pearson et al., 2009). Within a representative region of a cortical row containing three BB pairs, Poc5-GFP resided in assembling BBs and was not detected in mature BBs, marked by prominent Poc1-mCherry signal (Fig. 2A). Image averaging of Poc5-GFP and Poc1-mCherry signals across 58 BB pairs (representative BB pair in Fig. 2A) consistently detected TtPoc5 exclusively at the assembling BB (Fig. 2B). Notably, the timing of TtPoc5 incorporation into assembling BBs appeared to precede that of the slowly incorporated Poc1, since Poc5-GFP was found in assembling BBs devoid of Poc1-mCherry (arrows in Fig. 2A). Further, the absence of TtPoc5 in mature BBs may signify why Poc5-GFP was not observed in the oral apparatus BBs (Fig. 2A), which contain primarily mature BBs, as indicated by enriched Poc1-mCherry.

### Dynamic incorporation and removal of TtPoc5 in BBs

To verify that TtPoc5 localizes preferentially to assembling BBs, *Tetrahymena* cells with endogenously tagged Poc5-GFP were cultured in three conditions that elicit different cellular responses to BB assembly and maintenance: growth medium, starvation medium, and starvation medium followed by release into growth medium (Fig. 3A). During logarithmic growth, *Tetrahymena* BBs are stabilized for the entirety of the cell cycle and BB assembly is not synchronous, thus, the majority of cortical row BBs at any given time are mature with a minimal number of newly assembled BBs (Bayless et al., 2015; Galati et al., 2015). In growth, Poc5-GFP was only detected in a small proportion of cortical row BBs (arrowheads in Fig. 3A), which was consistent with TtPoc5 exclusively residing in assembling BBs (Fig. 2A,B). When *Tetrahymena* cells are shifted from growth medium into starvation medium, cells arrest in G1 preventing new BB assembly (Pearson et al., 2009). Under media starvation for 24 hours when only mature BBs persisted, Poc5-GFP signal was not detected in any cortical row BBs (Fig. 3A). Further, when starved cells are shifted back into growth medium this releases cells from cell cycle arrest (starve and release) and correspondingly initiates a synchronous wave of new BB assembly (Pearson et al., 2009). In contrast to the depleted Poc5-GFP signal in starved cells, there was a marked enrichment of Poc5-GFP-positive BBs during synchronized BB assembly in starved and released cells (arrowheads in Fig. 3A).

**Fig. 3.**
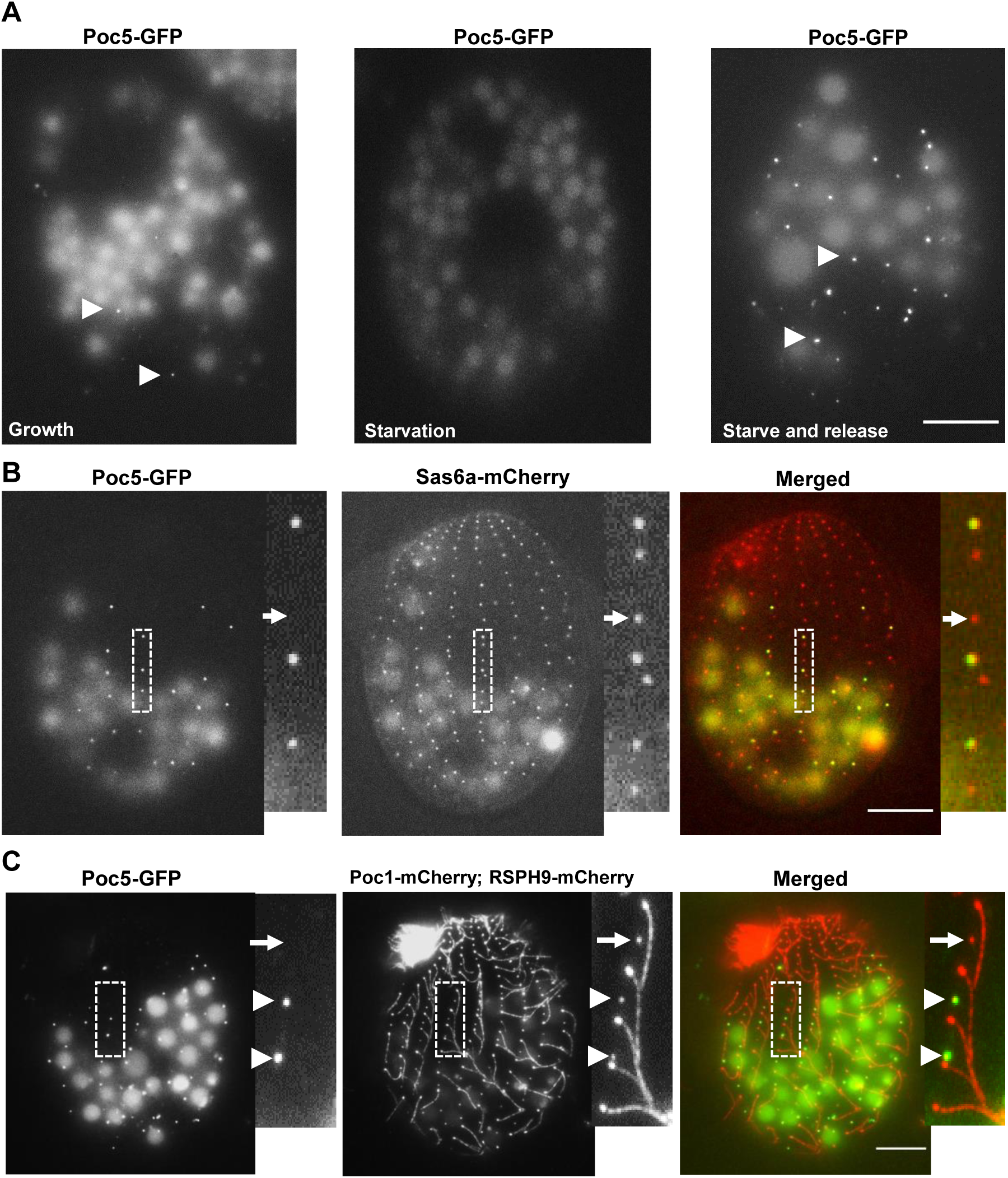
TtPoc5 is enriched during BB assembly and removed prior to cilia formation. **(A)** Live-cell imaging of endogenously tagged, C-terminal Poc5-GFP during logarithmic growth, in starvation, and after release back into growth medium (starve and release). During logarithmic growth, BB assembly occurs throughout the cell cycle and low levels of Poc5-GFP signal are detected at a single moment (indicated with arrowheads). In starvation, cells are arrested in G1 preventing new BB assembly and Poc5-GFP is not detectable. Overnight starvation followed by release into growth medium (stimulating new BB assembly) enriches the Poc5-GFP signal (marked with arrowheads). **(B,C)** Live-cell imaging of starved and released cells with representative sections of cortical rows (marked with dashed boxes) expanded in the smaller panels. **(B)** Endogenous Poc5-GFP is co-expressed with Sas6a-mCherry (early marker of BB cartwheel structure) to assess the timing of TtPoc5 BB incorporation. Sas6a-mCherry labels all BBs whereas Poc5-GFP is only at the anterior BB in a pair. Poc5-GFP signal always coincides with Sas6a-mCherry signal, therefore TtPoc5 BB incorporation does not precede that of Sas6a. **(C)** Endogenous Poc5-GFP is co-expressed with Poc1-mCherry (BB marker) and RSPH9-mCherry (evenly labels ciliary axonemes) to assess the timing of Poc5 removal from maturing BBs. Poc5-GFP removal precedes cilia formation since Poc5-GFP-positive BBs are not ciliated (indicated with arrowheads) and maturing, non-ciliated BBs with detectable Poc1-mCherry (highlighted with an arrow) can be devoid of Poc5-GFP signal. Scale bars: 10µm.

To better understand the timing of TtPoc5 BB incorporation relative to early assembly, cells co-expressing Poc5-GFP and Sas6a-mCherry were used since Sas6a is a critical component of the early-forming BB cartwheel structure that templates the nine-fold symmetry of BBs (Culver et al., 2009). In *Tetrahymena*, the cartwheel is a stable structure that persists from early BB assembly through maturation, therefore Sas6a-mCherry evenly labeled all cortical row BBs (Fig. 3B) (Bayless et al., 2015). To increase the number of Poc5-GFP-positive BBs for this analysis, cells were starved and released. Poc5-GFP localized exclusively to the assembling BBs within BB pairs and Poc5-GFP-positive BBs were always marked with Sas6a-mCherry (Fig. 3B). Given that Poc5-GFP-positive BBs were never devoid of Sas6a-mCherry, TtPoc5 BB incorporation preceded that of Poc1 (Fig. 2A) but did not precede early cartwheel formation (Fig. 3B). Notably, Sas6a-mCherry-positive BBs lacking any detectable Poc5-GFP were also observed (arrows in Fig. 3B), which are likely mature BBs where TtPoc5 has been removed.

To elucidate the timing of TtPoc5 BB removal relative to BB maturation and the onset of cilia formation, starved and released cells were used that co-expressed Poc5-GFP, Poc1-mCherry (BB marker), and a tagged *Tetrahymena* ortholog of the ciliary-specific protein, radial spoke head 9 (RSPH9-mCherry; gifted by Dr. Chad Pearson) (Yang et al., 2006). As seen in Fig. 3C, RSPH9-mCherry evenly labelled ciliary axonemes along cortical rows as well as in the mature oral apparatus, and there was no detected signal overlap with Poc5-GFP in BBs. Within a representative region of a cortical row (boxed area in Fig. 3C), Poc5-GFP colocalized with Poc1-mCherry in anteriorly-positioned BBs of pairs (arrowheads in Fig. 3C) but was not observed in mature BBs with prominent RSPH9-mCherry labelled cilia. The absence of TtPoc5 in ciliated BBs suggested that TtPoc5 removal from BBs was either prior to or prompted by the onset of cilia formation (arrowheads in Fig. 3C). In *Tetrahymena*, not all mature BBs are ciliated, providing an intermediate stage between BB maturation/stabilization and cilia formation for further dissection of this timing (Bayless et al., 2012; Nanney, 1975, 1971). As shown in Fig. 3C, mature nonciliated Poc1-mCherry-positive BBs were present that lacked Poc5-GFP signal (denoted with arrows), indicating that TtPoc5 removal occurred prior to the onset of cilia formation. Collectively, TtPoc5 is transiently incorporated into assembling BBs and is then removed prior to BB maturation, suggesting that TtPoc5 potentially functions during this dynamic stage of BB development and drawing similarities with the known function of hPOC5 in centriole elongation/maturation (Azimzadeh et al., 2009).

### Loss of TtPoc5 leads to overproduced cortical row BBs and a reduction of cilia

To determine the BB function(s) of TtPoc5, a complete genomic knockout (KO) strain was generated by replacing the *TtPOC5* open reading frame (ORF) through homologous recombination with a high efficiency, codon-optimized NEO2 cassette (coNEO2) built for usage in *Tetrahymena* and used for positive selection of transformants (Fig. 4A) (Mochizuki, 2008). The *TtPOC5* KO strain was validated by PCR using isolated genomic DNA from wild-type (WT) control and *poc5Δ* cells (Fig. 4B). RT-PCR using isolated RNA from WT and *poc5Δ* cells further confirmed that *poc5Δ* cells lacked *TtPOC5*, revealing expression of *coNEO2* only in *poc5Δ* cells and expression of *TtPOC5* only in WT cells (Fig. S1).

**Fig. 4.**
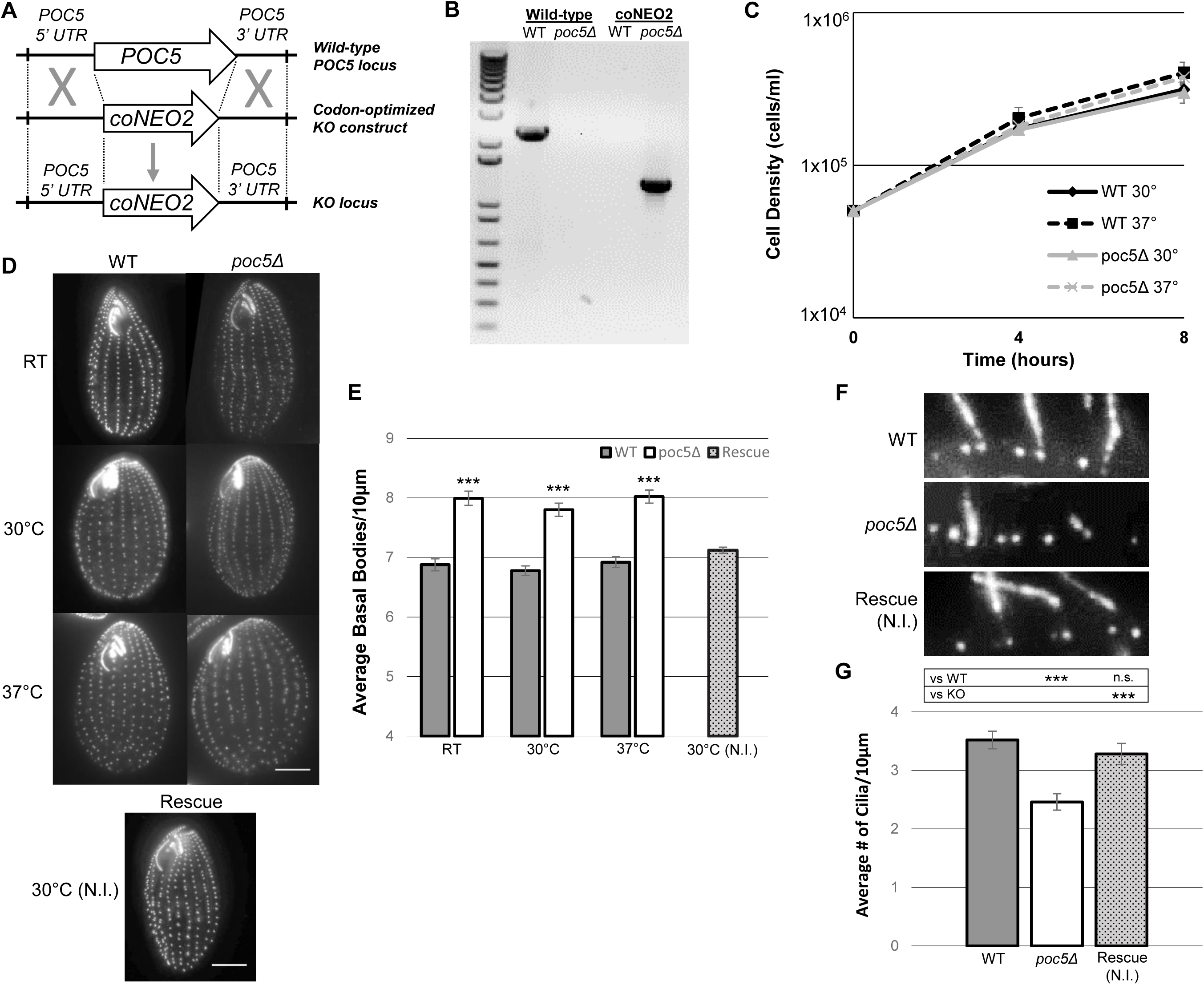
*poc5Δ* cells are viable and display overproduced BBs but fewer cilia. **(A)** Schematic outlining the homologous recombination strategy for generating a complete micronuclear knockout of *TtPOC5*, where the *TtPOC5 ORF* is replaced with a codon-optimized NEO2 (coNEO2) cassette for drug selection. **(B)** PCR confirming success of the knockout strategy, with only wild-type (WT) cells containing the *TtPOC5 ORF* and only *poc5Δ* cells containing the coNEO2 cassette. **(C)** Growth curves for WT and *poc5Δ* cells grown at either 30°C or 37°C for eight hours in SPP medium. Cell density (cells/ml) measurements are gathered at the initiation of the experiment (cultures starting at 0.5×10^5^ cells/ml), and four and eight hours after initiation. *n=3* analyzed samples per strain and incubation temperature. Error bars: SD. **(D)** Representative WT and *poc5Δ* cells stained with an antibody against Cen1 (labels BBs) after incubation at either room-temperature (RT), 30°C, or 37°C. Loss of TtPoc5 does not disrupt the organization/orientation of cortical row BBs. Bottom panel is representative *poc5Δ* rescue (Rescue) cell at 30°C with no CdCl_2_ induction (N.I.) due to leakiness of the *MTT1* promoter, where *TtPOC5* is reincorporated in *poc5Δ* cells through transformed *MTT1pr-GFP-Poc5*. Scale bars: 10µm. **(E)** BB density quantification (average BBs/10 µm) for WT and *poc5Δ* cells at RT, 30°C, and 37°C. Significant BB overproduction is observed in *poc5Δ* cells at all tested growth temperatures and rescued with MTT1pr-GFP-Poc5 to near WT levels at 30°C. *n=*300 total counts (5 counts per cell across 20 cells, in triplicate) per condition. Error bars: SEM. Student’s t-test calculation used to derive *P* values. ***, *P* < 0.001. **(F)** Representative 10 µm sections of cortical rows with an antibody against polyglutamylation (labels ciliary axonemes and BBs). **(G)** Ciliary density quantification (average # cilia/10 µm) for WT and *poc5Δ* cells at 30°C reveals significantly reduced ciliary density in *poc5Δ* cells that is rescued with *TtPOC5* reincorporation. *n=*100 total counts per condition. Error bars: SEM. Student’s t-test calculation used to derive *P* values. ***, *P* < 0.001.

*TtPOC5* is not essential for cell viability given that *poc5Δ* cells persist indefinitely; and through time-course analysis, *poc5Δ* cells exhibited similar growth rates to WT cells at both an optimal growth temperature for *Tetrahymena* (30°C) and at a restrictive temperature (37°C) used to identify temperature-sensitive mutants (Fig. 4C). From a gross morphological standpoint, *poc5Δ* cells grown at room-temperature (RT), 30°C, and 37°C displayed properly oriented BBs organized along cortical rows, as well as morphologically normal oral apparatuses (Fig. 4D). Since BB orientation was maintained in *poc5Δ* cells, the spatial organization of cortical row BBs in *poc5Δ* and WT cells was assessed by quantifying the BB density (average BBs/10 µm) along cortical rows (Fig. 4E). Measuring BB density in growing *Tetrahymena* cells is a powerful method for identifying disrupted BB homeostasis since the number of cortical row BBs is maintained at a nearly constant number (Frankel, 2008, 1980; Nanney, 1971, 1968, 1966). At all tested growth temperatures (RT, 30°C, 37°C), *poc5Δ* cells had a significantly greater cortical row BB density (8.0, 7.8, 8.0 BBs/10 µm, respectively) than WT cells (6.9, 6.8, 6.9 BBs/10 µm, respectively), suggesting that TtPoc5 potentially functions to inhibit BB overproduction or that loss of TtPoc5 elicits a compensatory cellular response leading to BB overproduction (Fig. 4E).

To determine that this observed BB overproduction was a direct result of loss of TtPoc5, a rescue strain was generated that reintroduced *TtPOC5* in *poc5Δ* cells through transformation of a cadmium-chloride (CdCl_2_)-inducible rescuing construct, *MTT1pr*-GFP-Poc5 (Fig. 4D). To assess rescue, the BB density was quantified in cells grown at 30°C with no added CdCl_2_ (no induction; N.I.), due to the established leakiness of the *MTT1* promoter (Heydeck et al., 2016; Shang et al., 2002) and because *MTT1pr-*mCherry-Poc5 exogenously expressed with N.I. in a WT background localized to cortical row BBs similarly to endogenous Poc5-GFP (Figs. 2A,S2). Further, CdCl_2_-induced *MTT1pr-*mCherry-Poc5 overexpression led to an aberrant TtPoc5 BB localization pattern and formation of elongated, Poc5-positive fibers observed previously with overexpression of Poc5 (arrows in Fig. S2) (Dantas et al., 2013). As seen in Fig. 4D and quantified in Fig. 4E, *poc5Δ* rescue cells grown at 30°C with N.I. exhibited a reduced BB density (7.1 BBs/10 µm) to near WT levels (6.8 BBs/10 µm) upon *TtPOC5* reincorporation, demonstrating that TtPoc5 has a direct function in modulating *Tetrahymena* BB production. Interestingly, despite the cortical row BB overproduction seen with loss of TtPoc5, *poc5Δ* cells had fewer ciliated BBs than WT cells (Fig. 4F,G). For this analysis, the ciliary density (average # of cilia/10 µm) was examined in WT, *poc5Δ*, and *poc5Δ* rescue cells (N.I.) grown at 30°C with an antibody against polyglutamylation, which marked microtubule glutamylation in both BBs and cilia (Bayless et al., 2016; Wolff et al., 1992). As shown in Fig. 4F and quantified in Fig. 4G, *poc5Δ* cells nucleated on average 2.5 cilia/10 µm along cortical rows compared with 3.5 cilia/10 µm in WT cells, and *TtPOC5* reincorporation in *poc5Δ* rescue cells grown at 30°C with N.I. led to a significantly increased ciliary density (3.3 cilia/10 µm) to near WT levels. Collectively, phenotypic analysis of *poc5Δ* cells elucidated a direct function for TtPoc5 in cortical row BB production and uncovered a diminished capacity in *poc5Δ* cells to maintain WT ciliary density levels despite BB overproduction.

### Loss of TtPoc5 and Sfr1 results in exacerbated BB overproduction and cell death

A potentially antagonistic role for TtPoc5 in modulating BB production is interestingly not unique for *Tetrahymena* centrin-binding proteins, where the previously characterized loss of Sfr1 led to a significantly greater BB density (7.9 BBs/10 µm) than in WT cells (6.98 BBs/10 µm) (Heydeck et al., 2016). Beyond the phenotypic overlap in *poc5Δ* and *sfr1Δ* cells, the BLAST search that identified TtPoc5 through sequence homology with hPOC5 (described in Fig. 1) identified Sfr1 as the second orthologous hit to hPOC5 (overall identities, 40/172 [23%]; positives, 74/172 [43%]). This shared sequence homology between hPOC5, TtPoc5, and Sfr1 primarily stemmed from a CBR organization that appears to be distinctive to Poc5 orthologs, with these proteins typically containing 3 CBRs organized as an isolated CBR and two CBRs in tandem (Fig. 1A) (Azimzadeh et al., 2009; Heydeck et al., 2016; Stemm-Wolf et al., 2013). Despite this shared CBR organization, the highly conserved Poc5 box that is a hallmark of Poc5 orthologs (Fig. 1D) is notably absent in Sfr1, therefore TtPoc5 was considered the primary *Tetrahymena* Poc5 ortholog (Azimzadeh et al., 2009; Heydeck et al., 2016; Stemm-Wolf et al., 2013).

Given the phenotypic overlap in *poc5Δ* and *sfr1Δ* single KO cells and potential functional redundancy between TtPoc5 and Sfr1, double KO cells lacking both *TtPOC5* and *SFR1* (*poc5Δ;sfr1Δ* cells) were generated using established methods (Fig. S3) (Hai et al., 2000; Heydeck et al., 2016). Unlike *poc5Δ* and *sfr1Δ* single KO cells that are viable and persist indefinitely, *poc5Δ;sfr1Δ* cells are inviable following completed drug selection at 30°C, suggesting that the overlapping BB function of TtPoc5 and Sfr1 was vital for *Tetrahymena* cell viability (Fig. 5A) (Heydeck et al., 2016). To test whether the underlying *poc5Δ;sfr1Δ* cell lethality was due to the unsuccessful transmission of the drug-resistance neomycin cassettes, cells containing a germline deletion of *TtPOC5* or *SFR1* only in the transcriptionally-silent micronuclei (heterokaryons) were mated with WT cells to generate a strain heterozygous for both *TtPOC5* and *SFR1*, resulting in viable cells after 48 hours post drug (Fig. 5A) (Cassidy-Hanley, 2012; Hai et al., 2000). Further, WT cells lacking drug-resistance cassettes were mated together to test the efficacy of drug selection, resulting in incomplete drug selection by 24 hours post drug but cell death by 48 hours in drug (Fig. 5A). Thus, *poc5Δ;sfr1Δ* cell death by 48 hours in drug was due to the combined loss of TtPoc5 and Sfr1, revealing an essential function for these *Tetrahymena* centrin-binding proteins that was obscured in *poc5Δ* and *sfr1Δ* single KO cells due to some functional redundancy (Heydeck et al., 2016).

**Fig. 5.**
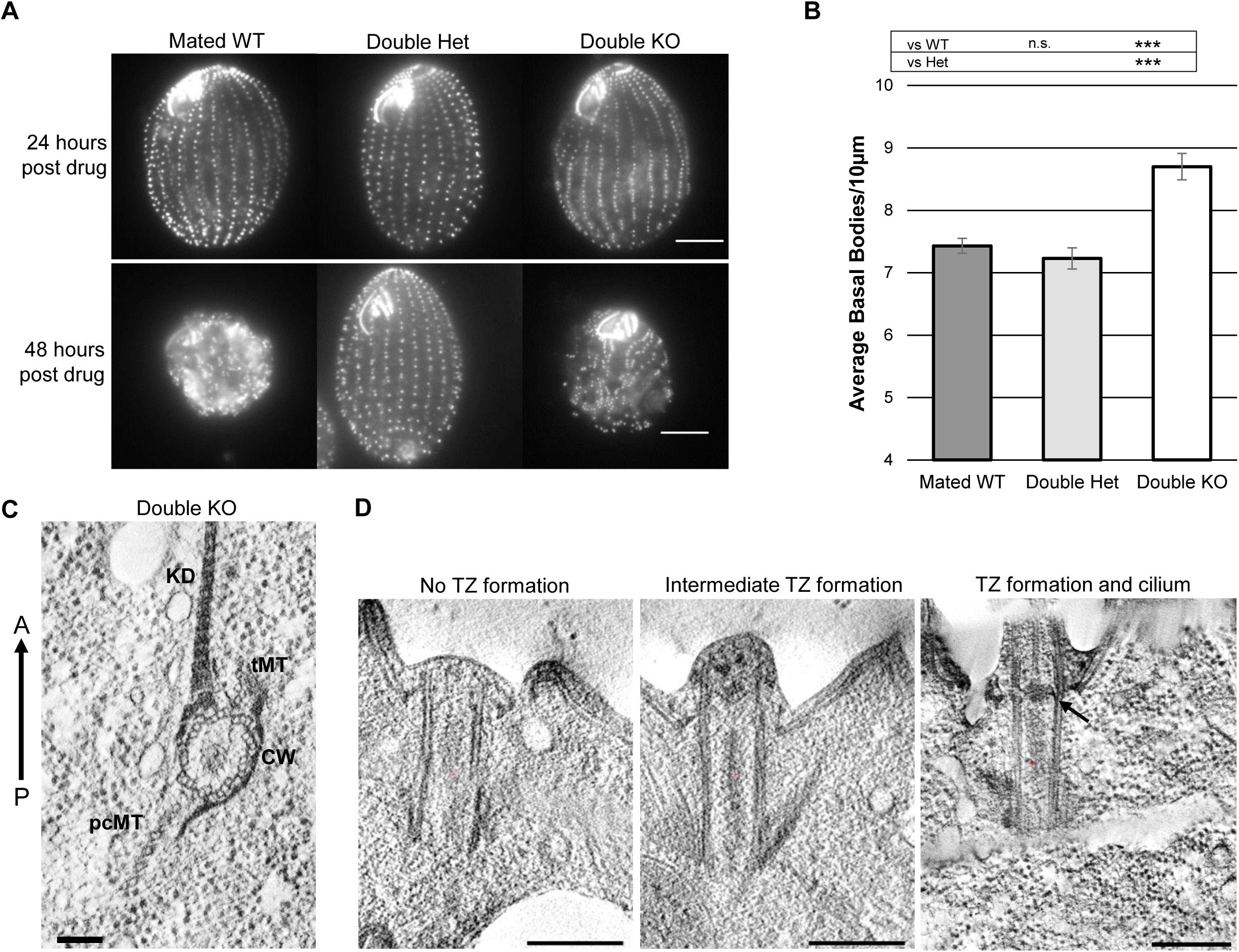
Loss of both TtPoc5 and Sfr1 results in cell lethality and exacerbated overproduction of defective BBs. A previously characterized centrin-binding protein, Sfr1, is the second best hPOC5 ortholog in *Tetrahymena* genome BLAST search, despite lacking the Poc5 box (Heydeck et al., 2016). Beyond sequence homology, Sfr1 has a similar role in modulating BB production, prompting the generation of double knockout (*poc5Δ;sfr1Δ*) cells using established methods (Hai et al., 2000). **(A)** Representative control mated WT, double heterozygous (het), and double knockout (KO) cells stained with anti-Cen1 antibody after 24 hours and 48 hours of drug selection following mating. Double KO cell death within 48 hours post drug selection is not due to a lack of drug resistance since double het. cells are viable after 48 hours in drug. Drug selection is effective since mated WT cells that do not carry drug resistance cassettes die after 48 hours in drug. Scale bars: 10µm. **(B)** BB density quantification (average BBs/10 µm) at 24 hours. Double KO cells have significantly higher BB density relative to mated WT and double het. cells. *n=*90 total counts (3 counts per cell across 15 cells, in 2 biological replicates). Error bars: SEM. Student’s t-test calculation used to derive *P* values. ***, *P* < 0.001. **(C,D)** Electron tomography of double KO BBs. **(C)** Cross-sectional view showing normal *Tetrahymena* BB accessory structures (kinetodesmal fiber (KD), transverse microtubules (tMT), post ciliary microtubules (pcMT)) and intact proximal cartwheel (CW). Scale bar: 100nm. **(D)** Longitudinal views showing BBs that do not template cilia and have varying degrees of transition zone (TZ) formation (left and middle panels). Right panel shows BB with a templated cilium and a morphologically normal TZ (arrow indicates electron dense acrosome). See Movie 1. Scale bars: 100nm.

To investigate the functional consequences of loss of both TtPoc5 and Sfr1 on BB production, phenotypic analysis of BB density was conducted after 24 hours in drug at 30°C when mated WT, double heterozygous, and *poc5Δ;sfr1Δ* cells persisted (Fig. 5A). Notably, mated WT cells were analyzed in parallel to control for BB rearrangement after mating and a potentially increased BB density from requisite BB assembly during cell division (Pearson and Winey, 2009). After 24 hours in drug, *poc5Δ;sfr1Δ* cells maintained properly oriented cortical row BBs and morphologically normal oral apparatuses, similar to what was observed in *poc5Δ* or *sfr1Δ* single KO cells (Fig. 4D) (Heydeck et al., 2016). As shown in Fig. 5A and quantified in Fig. 5B, the BB density in *poc5Δ;sfr1Δ* cells was significantly exacerbated (8.7 BBs/10 µm) compared with double heterozygous (7.2 BBs/10 µm) and mated WT cells (7.4 BBs/10 µm), while there was a no significant difference in BB density between double heterozygous and mated WT cells. Furthermore, the BB density in *poc5Δ;sfr1Δ* cells (8.7 BBs/10 µm) exceeded what was measured in *poc5Δ* cells grown at 30°C (7.8 BBs/10 µm) (Fig. 4E) and in *sfr1Δ* cells (7.9 BBs/10 µm) (Heydeck et al., 2016). Collectively, the increased cortical row BB overproduction in *poc5Δ;sfr1Δ* cells revealed functionally redundant roles for TtPoc5 and Sfr1 in BB production. Thus, *poc5Δ;sfr1Δ* cell death may be from an inability to compensate for a greatly increased level of BB production and consequently BB-dense cortical rows, which supports the importance of retaining a typical, highly regulated BB number in *Tetrahymena* (Frankel, 2008, 1980; Nanney, 1971, 1968, 1966).

Given that *poc5Δ;sfr1Δ* cells died by 48 hours in drug despite an abundance of BBs after 24 hours (Fig. 5A,B), an ultrastructural examination using electron microscopy (EM) was conducted to elucidate potential BB defects driving *poc5Δ;sfr1Δ* cell lethality (Fig. 5C,D). From cross-sectional views of *poc5Δ;sfr1Δ* cortical row BBs, it was apparent that the proximal cartwheel structures comprised of radially symmetric arrays of triplet microtubules were intact (Fig. 5C) (Allen, 1969; Bayless et al., 2015; Hirono, 2014; Kilburn et al., 2007). Additionally, BB accessory structures that aid in properly orienting and positioning BBs along cortical rows were also intact, which was not unexpected given the retained BB orientation in *poc5Δ;sfr1Δ* cells (Fig. 5C) (Bayless et al., 2015; Meehl et al., 2016, p. 1). Contrary to the proximal end of *poc5Δ;sfr1Δ* BBs, longitudinal views captured BBs docked at the cell surface with varying degrees of TZ formation at the distal end, suggesting that BB maturation was delayed or impaired upon loss of both TtPoc5 and Sfr1 (Fig. 5D, see Movie 1). Consequently, *poc5Δ;sfr1Δ* BBs with aberrant TZs did not template cilia at the cell surface (Fig. 5D) and lacked distinct TZ/axonemal features, including a transition from triplet-to-doublet microtubules (MTs), the electron dense axosome, and the central pair of MTs emanating from the axosome through the ciliary axoneme (Fig. S4, see Movies 2,3) (Bayless et al., 2015; Meehl et al., 2016; Sattler and Staehelin, 1974). Notably, BBs that templated cilia and exhibited morphologically normal TZs were also observed through EM (arrow marks axosome in Fig. 5D), which may be inherited BBs that formed prior to the loss of TtPoc5 and Sfr1 or the presence of these BBs could potentially indicate a dysregulated timing/delay of BB maturation in *poc5Δ;sfr1Δ* cells. By disrupting the timing of BB maturation and impairing TZ formation, the cortical row BB overproduction observed in *poc5Δ;sfr1Δ* cells was likely a compensatory response that resulted in an increased number of immature BBs, a diminished capacity to form cilia, and an inability to survive by 48 hours.

## DISCUSSION

In *Tetrahymena*, the tightly controlled timing and positioning of BB assembly, combined with the stabilization of BBs throughout the cell cycle, establishes a highly organized cytoskeleton with hundreds of BBs maintained along cortical rows and in the oral apparatus (Bayless et al., 2015). This intricately patterned and BB-rich cytoskeleton was previously exploited for characterizing the essential BB functions of centrin, which broadly encompass roles in BB assembly, orientation, separation, and stability (Kilburn et al., 2007; Alexander J Stemm-Wolf et al., 2005; Vonderfecht et al., 2011, 2012). Here, TtPoc5 is uniquely incorporated into assembling cortical row BBs and is then removed from mature BBs prior to the onset of cilia formation. *Tetrahymena* cells lacking *TtPOC5* are viable yet overproduce cortical row BBs and, despite this overproduction of BBs, exhibit a significantly reduced number of cilia along cortical rows. Upon reintroduction of TtPoc5 using an inducible promoter, the cortical row BB overproduction and reduced ciliary density seen with loss of TtPoc5 were both rescued. Interestingly, *Tetrahymena* has two *POC5*-like genes, *TtPOC5* and *SFR1*, that share homology within their CBRs and each encode BB-specific proteins with roles in modulating cortical row BB production (Heydeck et al., 2016). When *TtPOC5* and *SFR1* are co-deleted, *Tetrahymena* cells are no longer viable and display exacerbated levels of cortical row BB overproduction. Further, underlying BB maturation defects are apparent with varying degrees of distal TZ formation, leading to a consequently diminished capacity to nucleate a cilium. This study uncovers a specific requirement for Poc5 in building the distal portion of BBs, which corresponds with the distinct timing of TtPoc5 BB localization and may signify an added importance of centrin reported at the BB distal end (Kilburn et al., 2007). By revealing this functional redundancy, the dysregulation of BB number seen with loss of TtPoc5 and/or Sfr1 is likely a cellular response in *Tetrahymena* to compensate for delayed or defective BB maturation. Additionally, the essential role for TtPoc5 in *Tetrahymena* BB maturation provides a possible functional explanation for the observed association between *hPOC5* mutations and impaired cilia (Hassan et al., 2019; Oliazadeh et al., 2017; Patten et al., 2015; Weisz Hubshman et al., 2018).

### TtPoc5 is exclusively present in assembling BBs

In a previous analysis of hPOC5, this highly conserved centrin-binding protein was found to be recruited to procentrioles in the G2 phase of the cell cycle as procentrioles elongate and mature (Azimzadeh et al., 2009). Further, hPOC5 was observed in both the daughter and mother centrioles, similarly to centrin, indicating that once hPOC5 is recruited to procentrioles it is then stably incorporated. In comparison with hPOC5, this study finds endogenous TtPoc5 exclusively in assembling BBs after the initial cartwheel structure is formed and removal of TtPoc5 from mature BBs precedes ciliary assembly (Figs. 2, 3). This dynamic incorporation and removal of TtPoc5 relative to BB production has not been observed in previous analyses of other centrin-binding proteins, suggesting that the precise timing of TtPoc5 BB localization and function is important. Accordingly, overexpressing TtPoc5 using an inducible promoter appears to disrupt this critical timing, leading to the formation of elongated Poc5-positive fibers and an aberrant BB localization pattern with TtPoc5 residing in all cortical row BBs (Fig. S2). Interestingly, overexpression of Sfr1 did not produce Sfr1-positive fibers and Sfr1 was found to be stably present in all cortical row BBs as well as in the mature BBs within the oral apparatus (Heydeck et al., 2016). However, the functional significance and extent of this discrepancy in BB localization patterns is not currently understood primarily because of the technical challenges of endogenously tagging Sfr1, precluding a direct comparison with endogenous TtPoc5 (Heydeck et al., 2016). Given the stable presence of centrin in both mature BBs and centrioles, furthering our understanding of the molecular underpinnings driving Poc5 removal from mature BBs (and not mature centrioles), as well as illuminating the functional importance of Sfr1 residing in all BBs, will both be interesting to address in future research.

### Role of TtPoc5 in BB production and *Tetrahymena* BB number constancy

Beyond the established role for hPOC5 in centriole elongation/maturation, a Poc5 BB function has not been previously described (Azimzadeh et al., 2009; Chang et al., 2016; Chen et al., 2017; Comartin et al., 2013). In this study of TtPoc5 BB function, *poc5Δ* cells have a significantly increased cortical row BB density (Fig. 4E). This increase in BB number through BB overproduction is notable because *Tetrahymena* cells have an intrinsic capability to maintain a nearly constant total number of cortical row BBs, by altering the spatial organization of BBs through changes in the BB density and/or the overall number of cortical rows (Frankel, 2008, 1980; Nanney, 1971, 1968, 1966). The mechanism underlying the regulation of BB number constancy in *Tetrahymena* has not been elucidated, however, Sfr1 was similarly implicated in modulating cortical row BB production after observing more densely packed, supernumerary cortical rows in *sfr1Δ* cells (Heydeck et al., 2016). The absence of both TtPoc5 and Sfr1 leads to a greater increase in BB number (Fig. 5B), which has not been seen with genetic deletions of other characterized centrin-binding proteins, and uniquely implicates Poc5 regulating BB number. Notably, future studies are needed to address the significance of centrin-binding for Poc5 BB function, as well as the functional relevance of non-centrin interactors, including known ciliary and cytoskeleton components that bind hPOC5 (Hassan et al., 2019). Further, the highly conserved Poc5 box does not have a reported function, thus, this additional domain found only in Poc5 orthologs may specify Poc5 BB function.

Although most metazoan cell types lack the cortical organization of BBs that is characteristic of ciliates, there is a strong evolutionary conservation of molecules that tightly control both BB and centriole duplication irrespective of cortical organization (Firat-Karalar et al., 2014; Pearson and Winey, 2009; Rodrigues-Martins et al., 2008). Further, previous studies have established that centrosome amplification and the underlying deregulation of centriole number are widespread features of many human cancers (Chan, 2011; Marteil et al., 2018; Wong et al., 2015). Using a diverse panel of 60 human cancer cell lines derived from nine distinct tissues (the NCI-60 panel), significant centrosome amplification was evident in all tissue types and within half of the analyzed cell lines (Marteil et al., 2018). Further, there was marked variability in the percentage of cells with supernumerary centrioles, indicating that cancer cell lines possess distinct centriole numbers and levels of centrosome amplification. Interestingly, hPOC5 is capable of binding both human CETN2 and human CETN3, which have roles in promoting and antagonizing centrosome duplication, respectively (Sawant et al., 2015). Thus, depletion of CETN3 in HeLa cells results in a centriole overproduction, suggesting that centrins are important for modulating centriole number and this bidirectional contribution to centrosome duplication may be impaired in some human cancers. Collectively, these findings suggest that conserved mechanisms and/or signaling cues that are important for maintaining both centriole and BB number may exist, of which, Poc5 and centrins are good candidates for directing this process in metazoans.

### Requirement for Poc5 in BB maturation

The recruitment of hPOC5 to nascent centrioles occurs after initial assembly and coincides with building the centriole distal end, where hPOC5 and centrin colocalize (Azimzadeh et al., 2009; Paoletti et al., 1996b). Consistent with this localization pattern, hPOC5 is essential for centriole elongation/maturation but is not required for the initiation of procentriole assembly. In this study, cortical row BB production is significantly heightened in *Tetrahymena* cells lacking both *POC5* genes, and cross-sections of the BB ultrastructure reveal intact proximal cartwheel structures, indicating that Poc5 is not critical for initiating BB assembly (Fig. 5C). In contrast to the BB proximal end, cortical row BBs docked at or near the cell surface display variable TZ formation at the distal end, illuminating an important role for Poc5 in *Tetrahymena* BB maturation and a shared function for Poc5 in building the distal end of both BBs and centrioles (Fig. 5D) (Azimzadeh et al., 2009). Further, a role for Poc5 in BB maturation provides a functional explanation for the dynamic removal of TtPoc5 from mature BBs, suggesting that TtPoc5 is removed after completed formation of the BB distal end. Notably, a previous study in *Chlamydomonas* found concentrated centrin at the BB distal end and centrin-depletion uncovered a similar requirement for centrin in BB maturation, indicating that centrin and Poc5 may cooperate in this important process (Koblenz et al., 2003). Interestingly, centrin-depleted *Chlamydomonas* cells with BB maturation defects also contained aberrant BB numbers, including BB overproduction, suggesting that delayed and/or impaired BB development is functionally connected with dysregulation of BB number. Thus, the overproduction of cortical row BBs in *Tetrahymena* cells lacking TtPoc5 and Sfr1 is likely a mechanism to compensate for delayed and/or defective BB maturation rather than a novel role for Poc5 in antagonizing BB production. The functional extent of this compensatory mechanism requires further investigation, especially how BB production is impacted in response to distinct changes in the rate of BB biogenesis. Lastly, cancer cells frequently lose precise control over centriole length, and the recurrent feature of centriole size deregulation is associated with persistent centriole amplification (Marteil et al., 2018). Together, these findings may have uncovered a conserved cellular response that requires further examination, in which BBs and centrioles are overproduced when development of these structures, but not initial assembly, is compromised.

### Impact on ciliogenesis with loss of Poc5

Whereas hPOC5-depletion leads to impaired centriole maturation and subsequently defective cell cycle progression, the combined loss of TtPoc5 and Sfr1 in *Tetrahymena* results in immature cortical row BBs that consequently do not template cilia at the cell surface (Fig. 5D) (Azimzadeh et al., 2009). Additionally, structural features found within the distal TZ and ciliary axoneme, such as a triplet-to-doublet MT transition and a central MT pair, are not evident in cross-sections of the BB ultrastructure (Fig. S4) (Meehl et al., 2016). Taken together, this delayed and/or failed ciliary assembly is not unexpected since a mature TZ serves to compartmentalize the cilium and is required for ciliary assembly and maintained ciliary function (Czarnecki and Shah, 2012). Similarly, flagella are frequently absent in centrin-depleted *Chlamydomonas* cells as a result of delayed BB development and improper BB maturation (Koblenz et al., 2003). Notably, this role for Poc5 in BB maturation may provide a functional explanation for ciliary defects associated with *hPOC5* mutations, including shorter cilia observed with overexpression of a *hPOC5* variant associated with adolescent idiopathic scoliosis (Hassan et al., 2019; Patten et al., 2015; Weisz Hubshman et al., 2018; Xu et al., 2018). Also, Poc5 and centrin colocalize in the connecting cilium of photoreceptors, which corresponds structurally to the TZ and is vital for retinal function (Weisz Hubshman et al., 2018; Wheway et al., 2014; Ying et al., 2019). Although the function of Poc5 in the connecting cilium is not understood, a *hPOC5* mutation is associated with a form of retinal degeneration hallmarked by progressive loss of photoreceptors. Interestingly, a recent study of mouse centrins uncovered an important cooperative function between CETN2 and CETN3 in stabilizing the connecting cilium, suggesting that a role for Poc5 and centrins in BB maturation may translate to maturation of the analogous connecting cilium (Ying et al., 2019).

*Tetrahymena* cells are not viable when *TtPOC5* and *SFR1* are co-deleted, yet cells lacking either *POC5* gene persist indefinitely despite the presence of overproduced cortical row BBs. This suggests that the lethality observed in cells lacking both *POC5* genes is likely a direct consequence of delayed and/or failed ciliogenesis rather than an overproduction of BBs, given that the hundreds of cilia on the surface of *Tetrahymena* cells have essential roles for survival, including locomotory and feeding functions (Bayless et al., 2015; Gaertig et al., 2013). Interestingly, a co-occurrence of overproduced BBs and a reduced ciliary density is apparent in viable cells lacking TtPoc5 alone; indicating that further examination is needed to understand whether the degree in which cilia numbers are reduced, potentially in combination with exacerbated BB overproduction, underlie cell death when *TtPoc5* and *SFR1* are absent. Lastly, loss of TtPoc5 and Sfr1 may result in mis-localization of *Tetrahymena* Cen1 and/or Cen2 at the BB distal end, where proper localization and function of Cen1 is essential for cell viability (Alexander J Stemm-Wolf et al., 2005; Vonderfecht et al., 2012). Thus, increasing our understanding of the role of centrin in this process may be critical for understanding the full extent of Poc5 BB function. In summary, *Tetrahymena* has two paralogous *POC5* genes that are required for BB maturation, highlighting a requirement for Poc5 in building the distal end of both BBs and centrioles. In the absence of Poc5, *Tetrahymena* cells elicit a compensatory response to defective BB maturation that leads to BB overproduction and a consequential increase in the typically constant BB number. These abundant, immature BBs lack an ability to nucleate a cilium, illuminating the importance of Poc5 in ciliary assembly and potentially revealing the functional implications of *hPOC5* mutations on ciliary function.

## MATERIALS AND METHODS

### *Tetrahymena thermophila* strains and culture media

A WT strain derived from the progeny of a cross between *B2086* and *CU428* was used as the control for assaying the growth rate of the *poc5Δ* strain and for phenotypic analysis of the *poc5Δ* and *poc5Δ rescue* strains. A WT strain derived from the progeny of a cross between *CU427* and *SB1969* was the control comparison for phenotypic analysis of the *poc5Δ;sfr1Δ* strain (all parental WT strains from the Tetrahymena Stock Center, Cornell University, Ithaca, NY, USA). Cells were grown either at room-temperature (RT), 30°C, or 37°C as indicated, in 2% super-peptose (SPP) medium (Orias et al., 2000) to mid-log phase (~3×10^5^ cells/ml). Cell density measurements were determined by a Z2 Coulter Counter (Beckman Coulter, Brea, CA, USA). For starvation (arresting cells in G1), cells were grown to mid-log phase in SPP medium, washed in 10 mM Tris-HCl (pH 7.4), and then resuspended in 10 mM Tris-HCl (pH 7.4) for overnight incubation at 30°C. For starve and release experiments, cells were grown to mid-log phase in SPP medium, washed and resuspended in 10 mM Tris-HCl (pH 7.4) for overnight incubation at 30°C, and then released back into SPP medium for five hours (to stimulate synchronous BB assembly). For experiments with the cadmium-inducible *MTT1* promoter (Shang et al., 2002), cells were incubated overnight at 30°C in SPP medium either containing no cadmium chloride (CdCl_2_) or 0.5µg/ml CdCl_2_ at 30°C, as described.

### Identification of TtPoc5 and sequence comparison of Poc5 orthologs

*Tetrahymena* Poc5 (TtPoc5; TTHERM_00079160) was identified as the reciprocal best BLAST hit of full-length human POC5 (hPOC5)(Azimzadeh et al., 2009) in the *Tetrahymena* genome (Eisen et al., 2006). Of note, a prior evolutionary analysis of centriolar proteins used a previous alias of the same TtPoc5 ortholog, 5.m00513 (Hodges et al., 2010). The sequence alignment of hPOC5 and TtPoc5 was performed using ClustalW and the AA boxshading was performing using BoxShade, which are both available tools through the ExPASy Bioinformatics Resource Portal (https://www.expasy.org/genomics). Sequence logos were generated by WebLogo3 (Crooks et al., 2004; Schneider and Stephens, 1990) to visualize the AA composition of the conserved sequence motifs (canonical motif is Ax7LLx3F/Lx2WK/R) found within the three centrin-binding repeats (CBRs) of hPOC5 and TtPoc5 (Azimzadeh et al., 2009; Kilmartin, 2003). The 18 AA sequence motifs span positions 6-23, with five additional AAs upstream of the motifs (positions 1-5) included in the sequence logo. The size of the residue correlates with conservation and the color of the residue signifies the type of AA (hydrophobic is indicated in black, hydrophilic is blue, and neutral is green). The multiple sequence alignment of the highly conserved Poc5 box across select eukaryotes was performed using ClustalW and BoxShade. Select Poc5 orthologs as follows: human (NP_001092741), mouse (NP_080449), zebrafish (XP_691080), *Chlamydomonas reinhardtii* (Protein Id:9798, v4.0 through the Joint Genome Institute), and *Paramecium tetraurelia* (GSPATT00029936001, through *Paramecium* Genome Database).

### Plasmids and strain construction

The endogenous C-terminal GFP fusion used to localize TtPoc5 was made in the pGFP-LAP-NEO2 plasmid (based on the p4T2-1 vector) (Gaertig et al., 1994), which provides integration into the endogenous *TtPOC5* locus and expression of Poc5-GFP under control of the endogenous promoter. Briefly, 1kb of DNA from the 3’ end of *TtPOC5* (immediately upstream of the stop codon) was cloned adjacent to the GFP-LAP tag, and 1kb of 3’ sequence downstream of the stop codon was cloned into the vector flanking the NEO2 drug selection marker (conferring paromomycin resistance). A p4T21-MTT1pr-mCherry-Poc5 plasmid was created for TtPoc5 overexpression experiments, which provides integration into the endogenous *TtPOC5* locus and expression of the N-terminal mCherry-Poc5 fusion was under control of the cadmium-inducible *MTT1* promoter (Shang et al., 2002).

The *poc5Δ rescue* strain was generated by creating a *TtPOC5* rescue construct (pBS-*MTT1pr*-GFP-Poc5) that integrated into the *RPL29* locus and was under control of the cadmium-inducible *MTT1* promoter (Shang et al., 2002; Winey et al., 2012). This exogenous N-terminal GFP fusion was made by first cloning *TtPOC5* into the pENTR4 Dual Selection Entry Vector (Invitrogen, Carlsbad, CA, USA) and then using the Gateway cloning system (Invitrogen, Carlsbad, CA, USA), *TtPOC5* was subcloned into the pBS-*MTT1pr*-GFP-gtw vector (conferring cycloheximide resistance).

Sequence-confirmed constructs were linearized and transformed into the macronucleus of WT cells by biolistic bombardment using a PDS-1000 particle bombarder (Bio-Rad, Hercules, CA, USA) (Bruns and Cassidy-Hanley, 2000), except for pBS-MTT-GFP-POC5 (the poc5Δ rescue construct) which was transformed into the macronucleus of poc5Δ cells. For colocalization studies, cells expressing endogenous Poc5-GFP were transformed with endogenous Poc1-mCherry (Pearson et al., 2009), Sas6a-mCherry (Culver et al., 2009), and/or RSPH9-mCherry (gifted by Dr. Chad Pearson). In cells co-expressing Poc5-GFP, Poc1-mCherry, and RSPH9-mCherry, endogenous Poc1-mCherry was drug-selected with blasticidin while RSPH9-mCherry was drug-selected with cycloheximide (to allow for independent drug selection of each mCherry gene fusion).

### Generation of the *poc5Δ* and *poc5Δ;sfr1Δ* strains

The *poc5Δ* strain was generated by replacing the *TtPOC5* open reading frame with a high efficiency, codon-optimized *NEO2* (*coNEO2*) gene built for usage in *Tetrahymena* (Hai et al., 2000; Mochizuki, 2008). Targeted homologous recombination of the *TtPOC5* locus was achieved by using a knockout cassette with *coNEO2* flanked by 1kb of DNA upstream of the *TtPOC5* start codon and 1 kb of DNA downstream of the *TtPOC5* stop codon. For micronuclear transformation, mating strains *B2086/CrNeo* and *CU428/CrNeo* were used, which both contain a *NEO* gene in their macronucleus with two frameshift mutations to prevent DNA elimination of the selectable marker (Mochizuki et al., 2002; Yao et al., 2003). During conjugation, the *TtPOC5* knockout construct was initially transformed into the germline micronucleus using biolistic bombardment of DNA-coated gold particles, as previously described (Bruns and Cassidy-Hanley, 2000; Hai et al., 2000). Following biolistic bombardment, cells were incubated overnight in 10 mM Tris-HCl (pH 7.4) to finish conjugation, and then introduced to SPP medium for paromomycin drug-selection of positive transformants. Two different mating types of micronuclear knockout heterokaryons were generated through star crosses, as previously described (Hai et al., 2000). Single mating pairs were isolated from a mating between the two micronuclear knockout heterokaryon strains, and progeny were confirmed by PCR and RT-PCR for complete loss of *TtPOC5* (poc5Δ cells). For PCR validation: WT *TtPOC5* forward: ATGAATTCAAATAAGAATCAACCAAAGAAGAAA, WT *TtPOC5* reverse: TTTTTTGGTAGTTGTTGTTTTTGTTATTGC, *coNEO2* forward: ATTAATAACATTGCTGATGCTTTT and *coNEO2* reverse: GATTAATTACCTTCTAATAATTTGAAATAATTAATCC. For RT-PCR, RNA was isolated from WT and *poc5*Δ cells using the RNeasy Mini and QIAshredder kits (Qiagen, Hilden, Germany). cDNA was generated using the Superscript III One-Step RT-PCR system (Invitrogen, Carlsbad, CA, USA). Standard PCR followed: WT *TtPOC5* forward: ATGAATTCAAATAAGAATCAACCAAAGAAGAAA, WT *TtPOC5* reverse: TTTTTTGGTAGTTGTTGTTTTTGTTATTGC, *coNEO2* forward: ATTAATAACATTGCTGATGCTTTT, and *coNEO2* reverse: GAAGACGATAGAAGGCGATACG.

For generation of the *poc5Δ;sfr1Δ* strain, micronuclear knockout heterokaryon strains of two different mating types were initially generated with germline micronuclei homozygous for both *coNEO2* in the *TtPOC5* locus and NEO2 in the *SFR1* locus (Heydeck et al., 2016). Genotyping heterokaryons was performed by PCR since both cassettes conferred paromomycin drug resistance. PCR confirmation used: *coNEO2* forward: ATTAATAACATTGCTGATGCTTTT, *coNEO2* reverse: GATTAATTACCTTCTAATAATTTGAAATAATTAATCC, *NEO2* forward: AATCTACTAATTTGCTTTATTTTTCATAAGC, and *NEO2* reverse: TCCATACTTTGAAGATATCAAGC. An equal number of starved cells for each validated double heterokaryon strain were mixed together for mating overnight, and 24 hours after the initiation of mating an equal volume of 2x SPP medium was added. Cells were allowed to recover from mating in 2x SPP for four hours before adding paromomycin (200 µg/ml), this was considered the 0-hr timepoint for the *poc5Δ;sfr1Δ* double knockout (with basal body density measurements at 24 and 48 hours after paromomycin addition). In parallel to the double knockout mating, two control matings were performed as described with the double KO cells: WT x WT (*CU427 x SB1969*) and WT x heterokaryons (to generate cells that are heterozygous for *TtPOC5* and *SFR1*).

### Fluorescence Imaging

Images were collected using a Nikon Eclipse Ti inverted microscope (Nikon USA, Melville, NY, USA) with a CFI Plan Apo VC 60x H numerical aperture 1.4 objective and a charge-coupled device (CCD) CoolSNAP HQ2 camera (Teledyne Photometrics, Tuscon, AZ, USA). Metamorph Imaging Software (Molecular Devices, San Jose, CA, USA) was used for image acquisition, with all images acquired at RT.

For live-cell imaging, cells were grown in 2% SPP medium, washed in 10 mM Tris-HCl (pH 7.4), pelleted, and placed on microscope slides (VWR, Radnor, PA, USA). For immunofluorescence, cells were fixed using 3% formaldehyde and fixed cells were placed on poly-L-lysine coated multi-well slides (Polysciences Inc., Warrington, PA, USA). Cells were then blocked for one hour using phosphate-buffed saline (PBS) + 1% bovine serum albumin (BSA) before overnight incubation at 4°C in primary antibody, followed by a two-hour RT incubation in secondary antibody. All primary and secondary antibodies were diluted in PBS + 1% BSA. Primary antibodies used: α-TtCen1 (Alexander J. Stemm-Wolf et al., 2005) to label BBs and α-glutamylated tubulin (GT335 by AdipoGen, San Diego, CA, USA) to label ciliary axonemes (antibody also labels BBs) (Wolff et al., 1992). Secondary antibodies used: goat α-mouse or goat α-rabbit Alexa Fluor 488 (Invitrogen, Carlsbad, CA, USA). Cells were mounted in Citifluor mounting media (Ted Pella Inc., Redding, CA, USA).

In cells co-expressing Poc5-GFP and Poc1-mCherry, image averaging of GFP and mCherry signals was performed using ImageJ (National Institutes of Health, Bethesda, MD, USA) across 58 BB pairs. Images of Poc5-GFP and Poc1-mCherry BB signal were placed in a stack and the sum fluorescence of the stack was measured to create an averaged fluorescence image. The BB scaffolds were approximated using 200 nm diameter dashed circles, based on the average diameter of a *Tetrahymena* BB and the positioning of the center of the Poc1-mCherry signal.

### Electron Microscopy

Cells were prepared for electron tomography as described (Giddings et al., 2010; Meehl et al., 2009). Briefly, cells were gently spun into 15% dextran (molecular weight 9000–11,000; Sigma-Aldrich, St. Louis, MO, USA) with 5% BSA in 2% SPP medium. A small volume of the cell pellet was transferred to a sample holder and high-pressure frozen using a Wohlwend Compact 02 high pressure freezer (Technotrade International, Manchester, NH, USA). The frozen cells were freeze substituted in 0.25% glutaraldehyde and 0.1% uranyl acetate in acetone and then embedded in Lowicryl HM20 resin. Serial thick (250–300 nm) sections were cut using a Leica UCT ultramicrotome and section ribbons were collected on Formvar-coated copper slot grids. The grids were poststained with 2% aqueous uranyl acetate followed by Reynold’s lead citrate. 15 nm gold beads (BBI Solutions, Crumlin, UK) were affixed to both sides of the grid to serve as markers for subsequent tilt series alignment. Serial thin (80 nm) sections were collected and imaged to evaluate overall basal body structure and cortical row organization. Dual-axis tilt series of *Tetrahymena* cells were collected on a Tecnai F30 intermediate voltage electron microscope (Thermo Fisher Scientific, Waltham, MA, USA). Images were collected every one degree over a +/− 60 degree range using the SerialEM acquisition program (Mastronarde, 2005) with a Gatan Oneview camera at 1.2nm pixel. Serial section tomograms of *Tetrahymena* BBs were generated using the IMOD software package (Giddings et al., 2010; Kremer et al., 1996; Mastronarde, 1997). In total, five tomograms from the double KO and one serial tomogram from WT were reconstructed.

### Calculation of growth curves

Growth curves for the *poc5Δ* strain were calculated from cells grown in 2% SPP medium at 30°C and 37°C over the course of eight hours. Using a Z2 Coulter Counter (Beckman Coulter, Brea, CA, USA), cell density (cells/ml) measurements were taken at the 0-hour (with all cultures starting at 0.5×10^5^ cells/ml), 4-hour, and 8-hour timepoints. The final growth curves for WT and poc5Δ cells were based on averaging cell density measurements from three independent experiments. The *WT* strain used to calculate growth curves (derived from the progeny of B2086xCU428) was the same *WT* strain used for phenotypic analysis of the *poc5Δ* strain.

### Quantification of cortical row BB and ciliary density

The BB density (average BBs/10 µm) along cortical rows was quantified for WT and poc5Δ cells at RT, 30°C, and 37°C. BBs were labeled with the α-TtCen1 antibody and ImageJ (National Institutes of Health, Bethesda, MD, USA) was used to measure 10 µm regions along five separate cortical rows per cell (measurements on the side of the cell containing the oral apparatus and on the opposite side of the cell), across 20 cells in triplicate experiments. The final BB density was based on 300 total counts per condition. The same method was used to quantify the BB density for poc5Δ rescue and poc5Δ;sfr1Δ cells, but cells were only analyzed after growth in 2% SPP medium at 30°C. Quantification of the ciliary density (average cilia/10 µm) was quantified for WT, poc5Δ, and poc5Δ rescue cells at 30°C. Ciliary axonemes (and BBs) were labeled with the α-glutamylated tubulin (GT335) antibody and ImageJ was used to measure 10 µm regions on both sides of the cell (similar to measuring the BB density), with the average ciliary density based on 100 total counts per strain.

## ACKNOWLEDGEMENTS

We thank Chad Pearson for generously gifting the RSPH9-mCherry plasmid, Amy Fabritius for helpful discussions and reading of the manuscript, and Janet B. Meehl for her critical guidance of the electron microscopy work. Electron microscopy was performed at the Boulder Electron Microscopy Core Facility, University of Colorado, Boulder.

## AUTHOR CONTRIBUTIONS

W.H. executed all of the experiments with critical conceptualization from A.S.-W., except for the rescue and ciliary density experiments performed by B.A.B. and M.N., and the EM work performed by E.O.T. and C.O. M.W. was involved in experimental design and preparation of the manuscript.

---

**Fig. S1.**
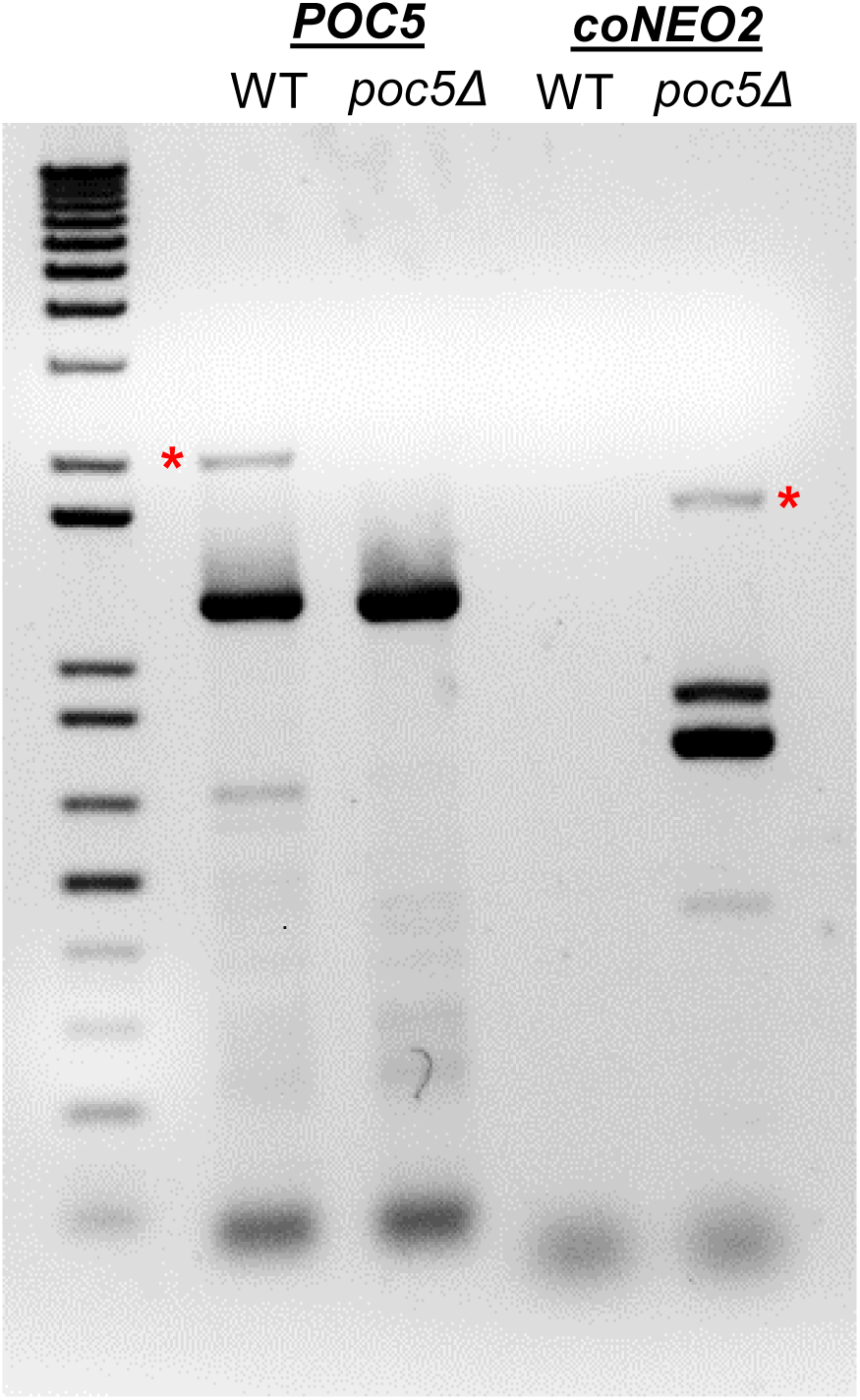
RT-PCR confirmation of *poc5Δ* cells. RT-PCR analysis using isolated RNA from WT and *poc5Δ* cells confirms *TtPOC5* KO and supplements PCR validation (Fig. 4B). *TtPOC5* expression is only observed in WT cells and *coNEO2* expression is only in *poc5Δ* cells. Amplified RT-PCR products specific to *TtPOC5* and *coNEO2* are indicated with red asterisks.

**Fig. S2.**
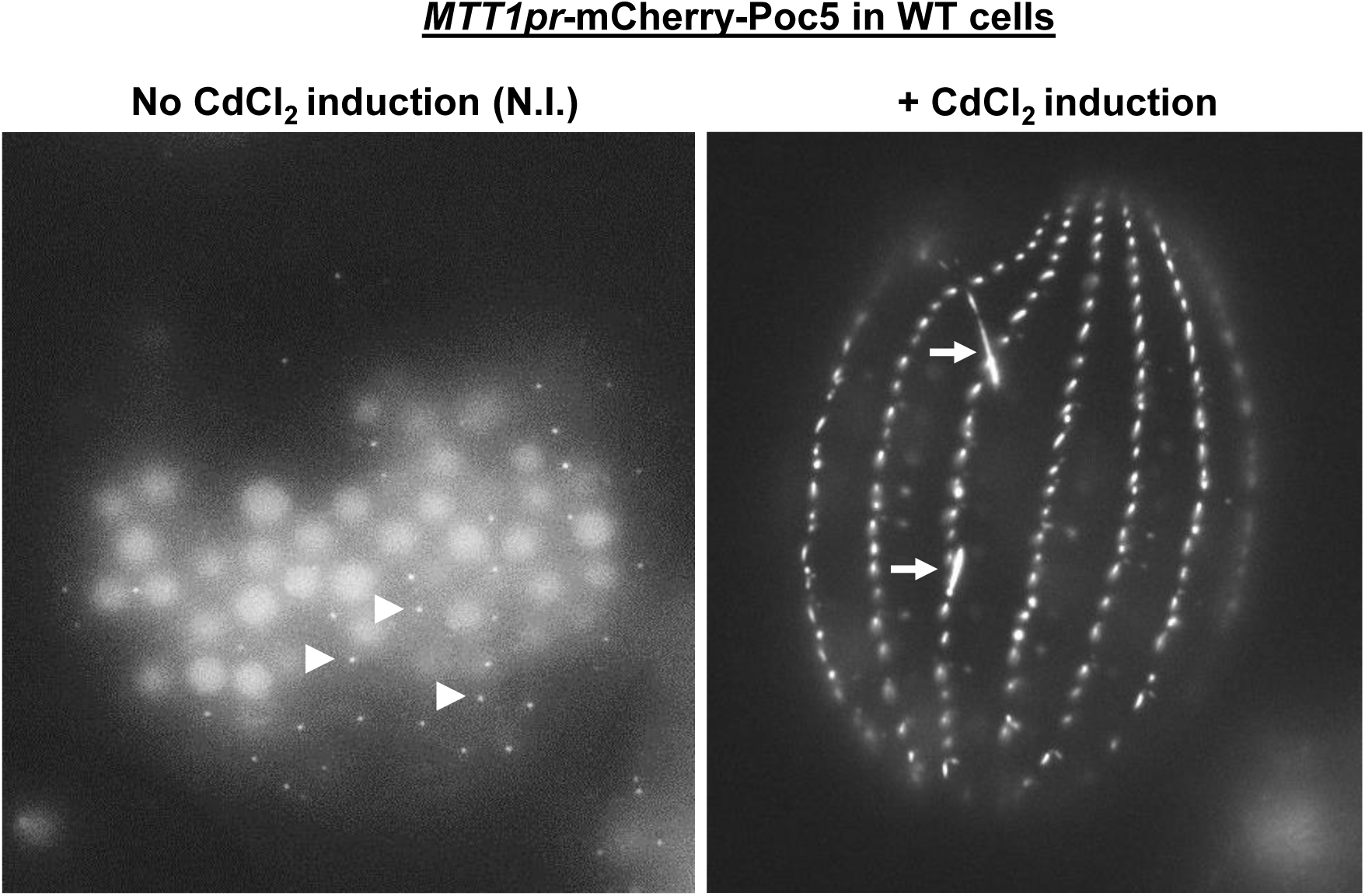
CdCl_2_-induced *MTT1pr*-mCherry-Poc5 overexpression leads to aberrant TtPoc5 BB localization. Due to leakiness of the *MTT1* promoter, exogenously expressed *MTT1pr*-mCherry-Poc5 BB localization (WT background) is assessed with both no CdCl_2_ induction (N.I.) and CdCl_2_-induced overexpression. *MTT1pr*-mCherry-Poc5 (N.I) localizes to a subset of cortical row BBs (marked with arrowheads), similar to endogenously expressed Poc5-GFP (Fig. 2A; Fig. 3A). In contrast, *MTT1pr*-mCherry-Poc5 (CdCl_2_ induction) appears to localize to all cortical row BBs yet remains absent in the oral apparatus. TtPoc5-positive fibers (labeled with arrows) are evident with CdCl_2_-induced *MTT1pr*-mCherry-Poc5 overexpression, as seen previously with Poc5 overexpression (Dantas et al., 2013).

**Fig. S3.**
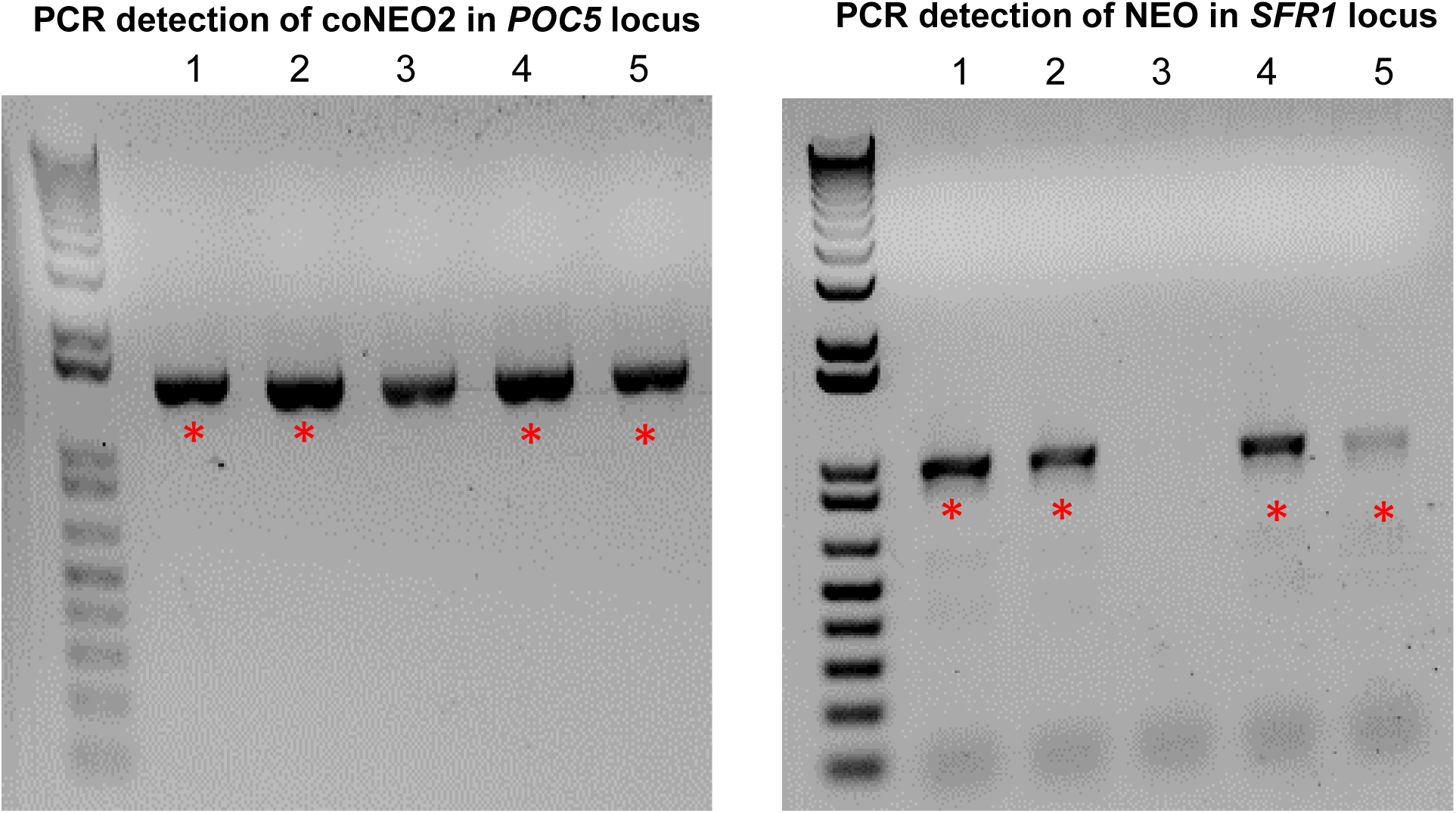
PCR confirmation of drug-resistant cells for generation of *poc5Δ;sfr1Δ* cells. Following paromomycin drug selection, PCR confirms cells with germline micronuclei containing codon-optimized NEO2 (coNEO2) in the *TtPOC5* locus and NEO2 in the *SFR1* locus (marked with asterisks). These cells were mated together to generate *poc5Δ;sfr1Δ* cells. Notably, one drug-resistant clone is identified (#3) that lacks NEO2 in the *SFR1* locus.

**Fig. S4.**
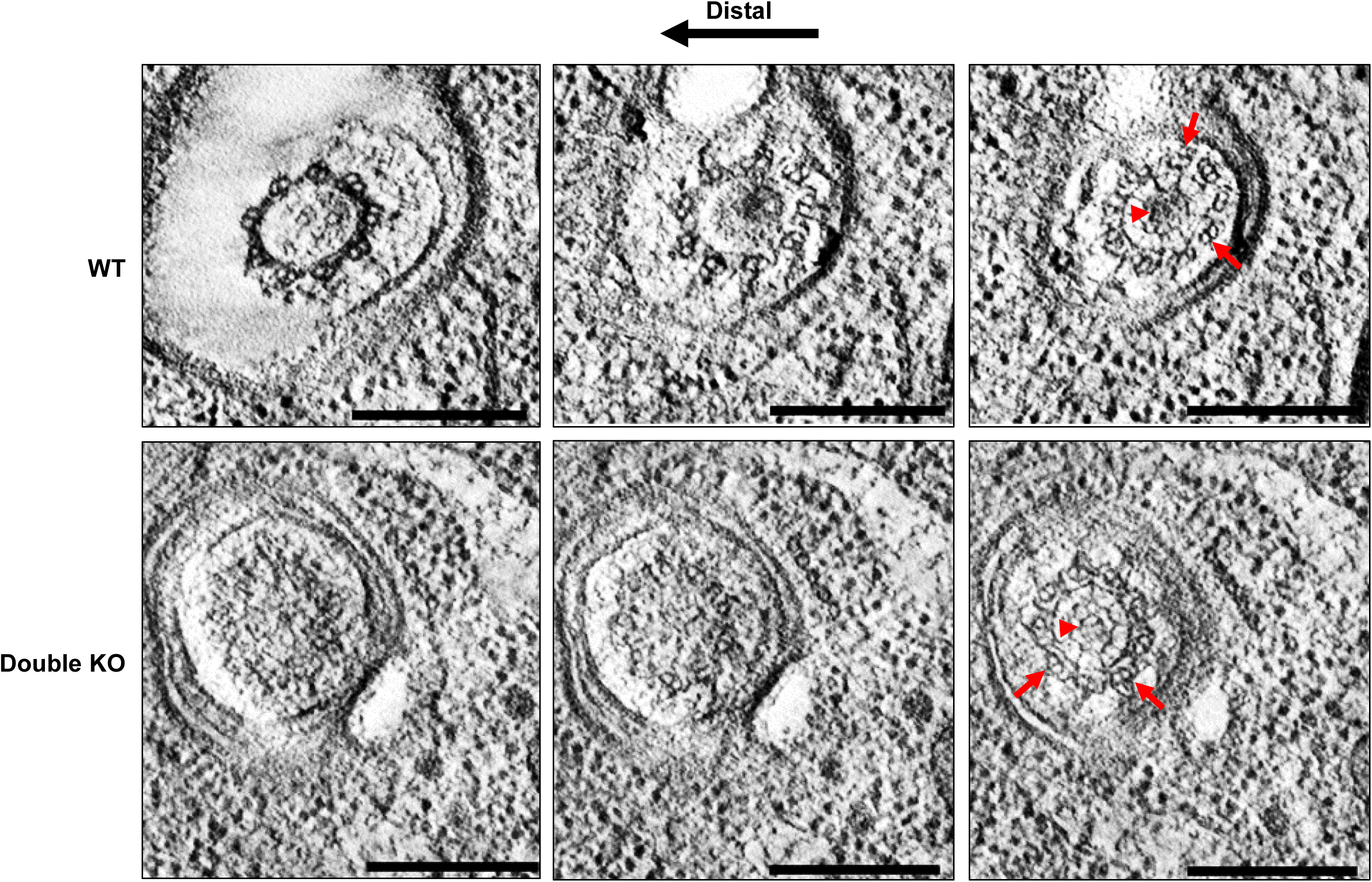
Ultrastructural BB maturation defects with loss of both TtPoc5 and Sfr1. Cross-sectional views from serial EM tomographic slices collected at the distal end of WT and double KO BBs. In double KO BBs, defective TZ formation is apparent, suggesting that loss of TtPoc5 and Sfr1 results in BB maturation defects. Double KO BBs lack characteristic TZ and ciliary axonemal features, including a typical transition from triplet-to-doublet microtubules (marked with arrows) and an electron-dense axosome containing a central microtubule pair (labelled with arrowheads). See Movies 2,3. Scale bars:100 nm.

**Movie 1. Double KO BBs have defective TZ formation.** Movie of serial EM tomographic slices through a portion of a double KO cell containing five BBs. Longitudinal view shows BBs with varying degrees of TZ formation. Still images from this volume presented in Fig. 5. Scale bar: 200nm.

**Movie 2. Serial EM tomographic slices through morphologically normal TZ.** Cross-sectional view of WT TZ with a typical transition from triplet-to-doublet microtubules, an electron dense axosome, and central pair microtubules at the distal end. Still images from this volume presented in Fig. S4. Scale bar: 200nm.

**Movie 3. Serial EM tomographic slices through defective TZ.** Cross-sectional view of a double KO TZ with apparently defective TZ formation and no detectable cilium. Still images from this volume presented in Fig. S4. Scale bar: 200nm.

